# Loss of Elp1 disrupts trigeminal ganglion neurodevelopment in a model of Familial Dysautonomia

**DOI:** 10.1101/2021.06.10.447739

**Authors:** Carrie E. Leonard, Frances Lefcort, Lisa A. Taneyhill

## Abstract

Familial Dysautonomia (FD) is a sensory and autonomic neuropathy caused by a mutation in *Elongator complex protein 1* (*ELP1*). FD patients have small trigeminal nerves and impaired perception of facial pain and temperature. These signals are relayed by nociceptive neurons in the trigeminal ganglion, a structure comprised of both neural crest- and placode-derived cells. Mice lacking *Elp1* in neural crest derivatives (“*Elp1 CKO*”) are born with smaller trigeminal ganglia, suggesting Elp1 is important for trigeminal ganglion development, yet the function of Elp1 in this context is unknown. We demonstrate Elp1 expression in both neural crest- and placode-derived trigeminal neurons, which our data suggest give rise to primarily TrkA- and TrkB/C-expressing neurons, respectively. While Elp1 is not required for initial trigeminal ganglion formation, *Elp1 CKO* trigeminal neurons exhibit abnormal axon outgrowth and decreased target innervation. Developing nociceptors that express the receptor TrkA are especially vulnerable to Elp1 loss. TrkA expression is decreased in *Elp1 CKO* trigeminal nerve endings, coinciding with increased cell death. Subsequently, fewer TrkA neurons are present in the *Elp1 CKO* trigeminal ganglion, indicating Elp1 supports the target innervation and survival of trigeminal nociceptors. These findings explain the loss of facial pain and temperature sensation in FD.

## Introduction

Hereditary sensory and autonomic neuropathies (HSANs) are a group of phenotypically similar, yet distinct, peripheral nervous system disorders that stem from unique mutations (Schwartzlow and Kazamel 2019). The most prevalent form is HSAN Type III, or Familial Dysautonomia (FD), which arises from an intronic mutation in the *Elongator Complex Protein 1 (ELP1)* gene (Anderson et al. 2001; Slaugenhaupt et al. 2001). This mutation causes tissue-specific mis-splicing of *ELP1*, subsequently reducing ELP1 protein in neurons (Slaugenhaupt et al. 2001; Hims et al. 2007). Elp1 is a requisite scaffolding protein of the six-subunit Elongator complex that functions in translation by modifying particular tRNAs (B. Huang, Johansson, and Byström 2005; B. Huang, Jian, and Byström 2008; Karlsborn, Tükenmez, Mahmud, et al. 2014; Xu et al. 2015; Esberg et al. 2006). Deletion of Elongator subunits has been shown to alter tRNA modifications and protein expression in several systems (C. Chen, Tuck, and Byström 2009; Esberg et al. 2006; Karlsborn, Tükenmez, Chen, et al. 2014; Goffena et al. 2018; Kojic et al. 2021). Consequently, loss or reduction of Elp1 protein leads directly and indirectly to the clinical phenotypes seen in FD, which include impaired pain and temperature sensation, feeding and swallowing difficulties, blood pressure instability, tachycardia, optic neuropathy, and gastrointestinal dysfunction, among other symptoms. The disease is fatal, with the majority of patients dying by age 50 (Gold-von Simson and Axelrod 2006). Autopsy reports and studies conducted in patients and animal models demonstrate FD phenotypes are caused by abnormal neurodevelopment, in addition to neurodegeneration across the lifespan of the affected individual (Pearson et al. 1978; Hunnicutt et al. 2012; George et al. 2013; Jackson et al. 2014; Li, Gruner, and Tourtellotte 2020; Won et al. 2019; Lefcort et al. 2017; Dietrich and Dragatsis 2016).

FD severely impacts the peripheral nervous system, composed of sensory and autonomic neurons that are mostly derived from neural crest cells and some cranial sensory neurons that arise from ectodermal placodes (Méndez-Maldonado et al. 2020; Steventon, Mayor, and Streit 2014; Kameneva and Adameyko 2019). Since mice that are null for *Elp1* are embryonic lethal (Y.-T. Chen et al. 2009; Dietrich et al. 2011), FD has been modeled in mice using conditional knockout of *Elp1* in neural crest cells (*Elp1^fl/fl^;Wnt1-Cre^+^*, abbreviated “*Elp1 CKO*”), which recapitulates several aspects of the human disease, including significant loss of sensory and autonomic neurons (George et al. 2013; Jackson et al. 2014). To date, these studies have focused on mechanisms of Elp1 in sensory and sympathetic neurons in the trunk. However, FD patients also experience cranial sensory deficits such as impaired sensation of facial pain and temperature, neurogenic dysphagia, absent corneal reflexes, and reduced basal lacrimation (Mendoza-Santiesteban et al. 2017; Palma, et al 2008; Barlow 2009; Geltzer et al. 1964; Gutiérrez, Norcliffe-Kaufmann, and Kaufmann 2015; Palma et al. 2018). These functions rely on input from the trigeminal nerve, the largest of the cranial nerves, which contains axons of sensory neurons that reside in the trigeminal ganglion. A quantitative MRI study revealed FD patients have smaller trigeminal nerves compared to healthy, age-matched controls, but there are no indications of progressive trigeminal nerve degeneration (Won et al. 2019). Moreover, *Elp1 CKO* mice are born with smaller trigeminal ganglia compared to controls (Jackson et al. 2014). Together, these findings suggest Elp1 may play an important role in the development of trigeminal sensory neurons, a critical aspect of FD that has yet to be explored.

While peripheral neurons in the trunk are exclusively neural crest-derived, cranial sensory neurons arise from two progenitor populations, neural crest cells and ectodermal placodes, which typically contribute neurons to spatially distinct regions of the cranial ganglia (York, Yuan, and McCauley 2020; Moody and Lamantia 2015; Park and Saint-Jeannet 2010; Steventon, Mayor, and Streit 2014). The trigeminal ganglion is unique in that it contains intermixed neural crest- and placode-derived neurons that rely on one another for proper migration, coalescence, and function during development (Blentic et al. 2011; D’amico-Martel 1982; D’Amico-Martel and Noden 1983; Hamburger 1961; Saint-Jeannet and Moody 2014; Shiau et al. 2008; Steventon, Mayor, and Streit 2014). Shortly after differentiation, trigeminal ganglion neuron subtypes are discernable by mutually exclusive expression of Tropomyosin receptor kinase (Trk) receptors, TrkA, TrkB, or TrkC, which are required for target innervation and long-term survival (E. J. Huang, Wilkinson, et al. 1999; Wilkinson et al. 1996; Scott-Solomon and Kuruvilla 2018; A. M. Davies 1997; Reichardt 2006; E. J. Huang, Zang, et al. 1999). Importantly, Trk expression generally correlates with the ultimate sensory modality encoded by a particular neuron; for example, small-diameter TrkA neurons are typically associated with pain and temperature perception (Mu et al. 1993; Genç, Ulupinar, and Erzurumlu 2005; d’Amico-Martel and Noden 1980; D’Amico-Martel and Noden 1983; A. Davies and Lumsden 1984). In dorsal root (sensory) and sympathetic (autonomic) ganglia, Elp1 is required for the generation and/or survival of TrkA and TrkB neurons, while TrkC neurons are spared during development (Jackson et al. 2014; George et al. 2013). In these contexts, neuronal loss has been attributed to decreased neurogenesis resulting from early differentiation and apoptosis of progenitors (George et al. 2013; Goffena et al. 2018), and to severe target innervation defects and neuronal apoptosis due to insufficient neurotrophic support (Jackson et al. 2014; Li, Gruner, and Tourtellotte 2020). Notably, other less common forms of HSANs arise from germline mutations in the genes encoding either *TrkA* or its high-affinity ligand, *nerve growth factor* (*NGF*) (Schwartzlow and Kazamel 2019). A growing body of evidence from animal models suggests Elp1 regulates TrkA signaling as well (Abashidze et al. 2014; Lefler et al. 2015; Naftelberg et al. 2016; Li, Gruner, and Tourtellotte 2020), although the relationship between Elp1 and TrkA has not been investigated in the cranial ganglia. While there are clear trigeminal sensory deficits in FD, the function of Elp1 in the trigeminal ganglion and its nerves remains unexamined. Moreover, it is unknown whether certain neuronal subtypes within the trigeminal ganglion are more vulnerable to Elp1 loss than others, or whether trigeminal sensory phenotypes in FD patients arise from defects in neural crest-derived neurons, placodal neurons, or both.

Here, we describe the first comprehensive analysis of neurodevelopmental changes in the trigeminal ganglion using an established *Elp1 CKO* mouse model of FD, where *Elp1* is deleted from neural crest cells and their derivatives (George et al. 2013). We demonstrate that Elp1 protein is enriched in the cytoplasm of differentiated neural crest- and placode-derived neurons in the trigeminal ganglion. While initial formation of the trigeminal ganglion is unaltered in *Elp1 CKO* mice, axon outgrowth, nerve formation, and nerve branching are severely diminished as development proceeds. These changes are accompanied by a drastic decrease in TrkA levels in the peripheral and central projections of trigeminal ganglion neurons as well as a significant reduction in the number of TrkA neurons within the ganglion. Finally, we demonstrate this loss of TrkA neurons due to neural crest-specific deletion of *Elp1* is correlated with increased cell death. Collectively, these findings indicate Elp1 is required for proper target innervation and survival of TrkA neurons in the trigeminal ganglion. Our results complement previous reports that demonstrate Elp1 modulates TrkA signaling, yet we now provide additional insight into nuanced phenotypes that may arise due to differences between nerve branches of the same ganglion. Moreover, by using a neural crest-specific knockout in the trigeminal ganglion, which contains neurons of both neural crest and placodal origin, we shed light on the neurodevelopmental dynamics and lineage of specific neuronal subpopulations in the trigeminal ganglion. Importantly, our findings introduce a cellular mechanism to explain the developmental defects occurring in the trigeminal ganglion, which ultimately contribute to the etiology of facial sensory deficits experienced by patients with FD.

## Results

### Elp1 protein is enriched in the cytoplasm of trigeminal ganglion sensory neurons

While ubiquitous Elp1 expression in rodent embryos has been previously reported (Mezey et al. 2003; George et al. 2013), the spatio-temporal distribution of Elp1 in the craniofacial complex has not been evaluated. To initially address this, we examined the *Elp1:LacZ* (previously “*Ikbkap:LacZ”*) reporter mouse that expresses β-galactosidase from the *Elp1* locus (George et al. 2013). Whole-mount preparations revealed widespread β-galactosidase staining in the head at embryonic day 10.5 (E10.5), with prominent expression in the neural tube, regions of the developing cranial ganglia, the pharyngeal arches, and the facial prominences (Fig. 1A). Horizontal sections through the head at E10.5 revealed diffuse β-galactosidase expression in the trigeminal ganglion, which, at this stage, is a newly condensed structure containing differentiated trigeminal placode-derived neurons and undifferentiated cranial neural crest cells (Fig. 1B, C, Karpinski et al.

**Figure 1.**
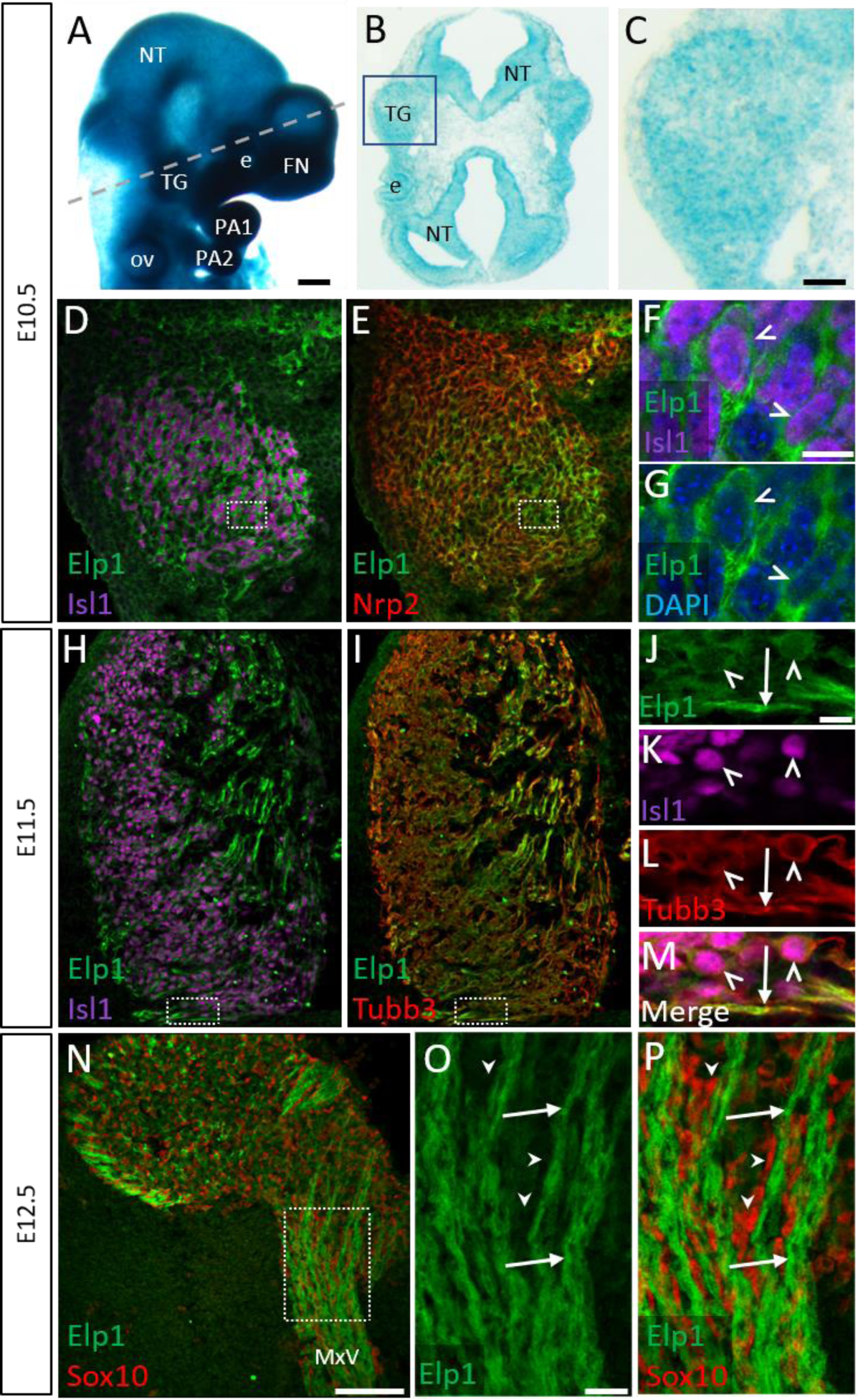
Elp1 protein is enriched in the cytoplasm of developing trigeminal ganglion neurons. (**A**) Lateral view of E10.5 *Elp1:LacZ* reporter mouse stained for β- galactosidase. Dashed line indicates the plane of section for the same embryo shown in **B**. (**B-C**) Representative horizontal section through embryo in **A** to reveal *Elp1* gene expression. Boxed region in **B** is magnified and shown in **C**. (**D-P**) Representative horizontal sections taken from wildtype E10.5 (**D-G**), E11.5 (**H-M**), or E12.5 (**N-P**) mouse embryos followed by fluorescent immunohistochemistry for Elp1 (**D-P**, green), Islet1 (**D**, **F**, **H**, **K**, “Isl1”, purple), Neuropilin2 (**E**, “Nrp2”, red), β-tubulin III (**I**, **L**, “Tubb3”, red), and Sox10 (**N**, **P**, red). Boxed region in **D** and **E** is magnified in **F-G** and shows Elp1 (green), Isl1 (purple), and DAPI-stained nuclei (blue). (**J-M**) Higher magnification of box in **H** and **I**. (**O**-**P**) Higher magnification of box in **N**. Carets indicate Isl1-positive and/or Tubb3-positive neuronal cell bodies (**F**, **G, J-M**). Arrows identify axons (**J**, **L, M, O, P**), while arrowheads point to Sox10-positive glial progenitors (**O-P**). Abbreviations: e – eye, FN – frontonasal prominence, NT – neural tube, ov – otic vesicle, PA – pharyngeal arch, TG – trigeminal ganglion. Scale bars: 400µm (**A**), also applies to **B**; 50µm (**C**), applies to **D, E, H, I**; 100µm (**N**); 10µm (**F**), applies to **G**; 10µm (**J**), applies to **K, L, M, O, P**.

2016). β-galactosidase expression was also apparent in the neural tube, the eye, and scattered throughout the cranial mesenchyme (Fig. 1B, C). Together, these results indicate the *Elp1* gene is expressed in cranial neural tissues, similar to findings in the trunk, and more specifically that *Elp1* is expressed in the newly formed trigeminal ganglion.

Interestingly, detection of Elp1 via immunohistochemistry revealed an overlapping, but more specific, expression pattern. At E10.5, Elp1 protein was abundant in the cytoplasm of differentiated neurons identified by expression of the transcription factor Islet1, and was not detected in the nucleus (Fig. 1D-G and Fig. 1–figure supplement 1A- D). Since the vast majority of neurons in the trigeminal ganglion are placode-derived at this stage, while surrounding neural crest cells have yet to differentiate (Karpinski et al. 2016), we conclude that Elp1 is initially expressed in trigeminal placode-derived neurons. In contrast, condensed neural crest cells within the trigeminal ganglion, identified by expression of the surface receptor Neuropilin2 or the transcription factor Pax3, and devoid of Islet1, expressed little to no Elp1 protein (Fig. 1E and Fig.1–figure supplement 1A-D). Compared to the diffuse *Elp1:LacZ* signal in the trigeminal ganglion at the same stage (Fig. 1A-C), this suggests *Elp1* may be post-transcriptionally regulated, leading to robust Elp1 protein expression in neurons upon differentiation.

**Figure 1–figure supplement 1.**
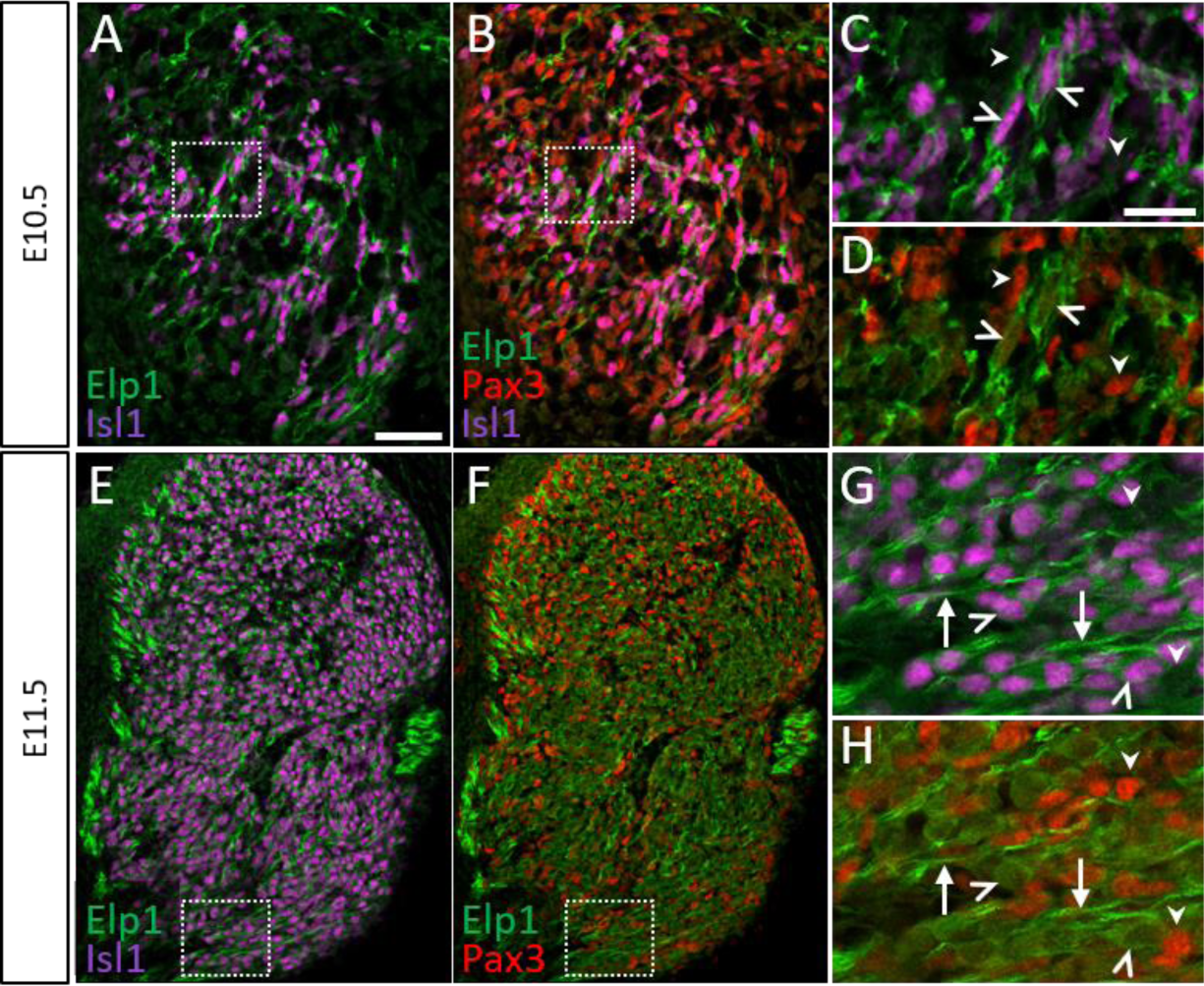
Elp1 is not expressed in Pax3-positive neural crest cells or glial progenitors in the developing trigeminal ganglion. (**A-H**) Fluorescent immunohistochemistry on representative horizontal sections from E10.5 (**A-D**) and E11.5 (**E-H**) wildtype mouse embryos demonstrating expression of Elp1 (**A-H**, green), Isl1 (**A-C**, **E**, **G**, purple) and Pax3 (**B**, **D**, **F**, **H**, red). Carets point to Isl1-positive neuron cell bodies (**C**, **D**, **G**, **H**). Arrows indicate axons (**G**, **H**), while arrowheads denote Pax3-positive neural crest cells (**C**, **D**) or glial progenitors (**G**, **H**). Scale bars: 50µm (**A**), applies to **B, E, F**; 20µm (**C**), applies to **D, G, H**.

At E11.5, when neural crest-derived neurons are now present in the trigeminal ganglion, Elp1 protein was observed in all differentiated neurons expressing β-tubulin III (Tubb3) and Islet1, irrespective of their developmental origin, with the highest signal detected in axons (Fig. 1H-M and Fig.1–figure supplement 1E-H). These findings are similar to previously reported observations in other peripheral ganglia (Abashidze et al. 2014; Hunnicutt et al. 2012; George et al. 2013). By E12.5, enrichment of Elp1 protein in the cytoplasm, especially axons, of Tubb3-positive neurons is still apparent, while Elp1 is absent or expressed at relatively low levels in neural crest-derived Sox10-positive glia within the ganglion and along the embryonic nerves (Fig. 1K-M). Collectively, these results suggest important functions for Elp1 in developing sensory neurons of the trigeminal ganglia.

### Neural crest-specific deletion of *Elp1* causes progressive morphological abnormalities within the trigeminal ganglion and its nerve branches

To understand the requirements for Elp1 during trigeminal ganglion development and to gain insight into the etiology of FD phenotypes associated with trigeminal ganglion dysfunction, an established mouse model was used, in which *Elp1* is conditionally deleted from neural crest cells and their derivatives via Cre-mediated recombination under the Wnt1 promoter (*Elp ^flox/flox^; Wnt1-cre^+^*, “*Elp1 CKO*”) (George et al. 2013). The anatomy and morphology of the embryonic trigeminal ganglion and associated nerves of *Elp1 CKO* and littermate controls were evaluated over developmental time. The normal progression of trigeminal nerve development between E10.5 and E13.5 is depicted in Figure 2, while *Elp1 CKO* trigeminal ganglion and nerve phenotypes relative to controls are summarized in Table 1.

**Figure 2.**
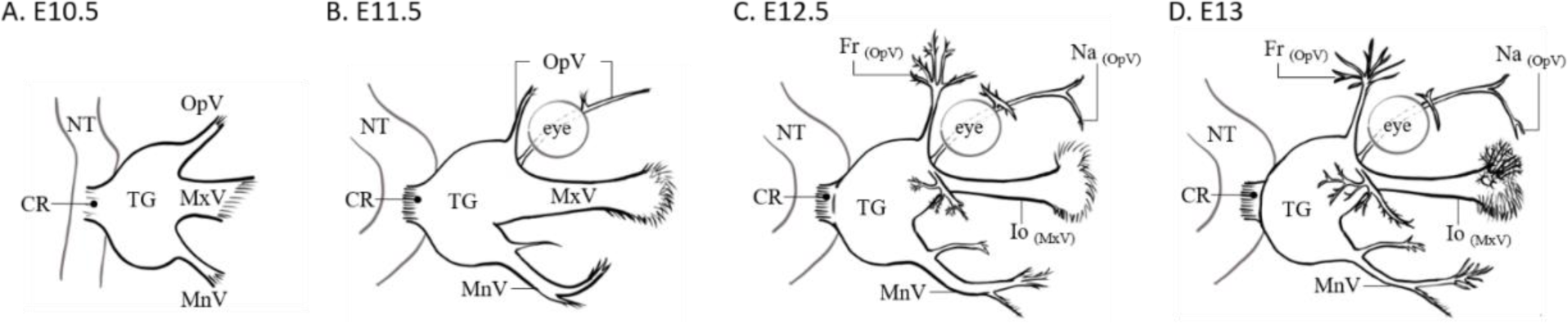
Schematic of normal mouse trigeminal ganglion and nerve anatomy over developmental time. (**A-D**) Diagram labels relevant ganglion and nerve anatomy in E10.5 (**A**), E11.5 (**B**), E12.5 (**C**), and E13 (**D**) mouse embryos for reference in Figures 3-6. Abbreviations: CR – central root, Fr – frontal nerve, Io – infraorbital nerve, MnV – mandibular nerve, MxV – maxillary nerve, Na – nasal nerve, NT – neural tube, OpV – ophthalmic nerve, TG – trigeminal ganglion.

**Table 1.** Comparison of Elp1 CKO Trigeminal Ganglion and Nerve Phenotypes Relative to Control Across Developmental Time. Summary of observations of the trigeminal ganglion, ophthalmic nerve, maxillary nerve, mandibular nerve, and central nerve root in *Elp1 CKO* between E10.5 and E13.5, as compared to control littermates. *, mandibular nerve was difficult to visualize at these stages, so no observations were recorded.

|  | Gestational Age |  |  |  |
| --- | --- | --- | --- | --- |
|  | E10.5 | E11.5 | E12.5 | E13-13.5 |
| **Ganglion** | No difference; Tubb3-positive placodal neurons and Sox10-positive neural crest cells are present/ distributed throughout forming ganglion | Slightly less compact | Less compact; increased cell death | Less compact; TrkA expression present but reduced; fewer TrkA-positive neurons present |
| **Ophthalmic N.** | Present; no difference in length | No difference | Axons straying from established fascicles; decreased branching complexity; nasal nerve absent | Decreased nerve complexity even more apparent; “dying back” of axons; nasal nerve still absent; reduction in TrkA expression |
| **Maxillary N.** | Present; no difference in length | Disorganized; many axons stray from established fascicles as they traverse toward the whisker pad | No change in whisker pad innervation | Decreased innervation of whisker pad; reduction in TrkA expression |
| **Mandibular N.** | Present; no difference in length | Disorganized and less compact; "hairy" appearance due to straying axons | * | * |
| **Central root axons** | No difference | Less compact | Smaller | Smaller; reduction in TrkA expression |

When compared to littermate controls at E10.5, initial formation of the trigeminal ganglion appeared normal in *Elp1 CKO*, as demonstrated by whole-mount Tubb3 immunohistochemistry, which labels all neurons (Fig. 3A, B). Gross anatomy was intact, including a condensed ganglion and the emergence of the ophthalmic, maxillary, and mandibular nerve branches (Fig. 3A, B). Immunohistochemistry on E10.5 tissue sections revealed both Tubb3-positive placodal neurons and Sox10-positive undifferentiated neural crest cells were present and appropriately distributed throughout the forming ganglion (Fig. 3C, D). Altogether, these data suggest Elp1 is not required for cranial neural crest cells to migrate to the trigeminal anlage and coalesce with placode-derived neurons for ganglion assembly.

**Figure 3.**
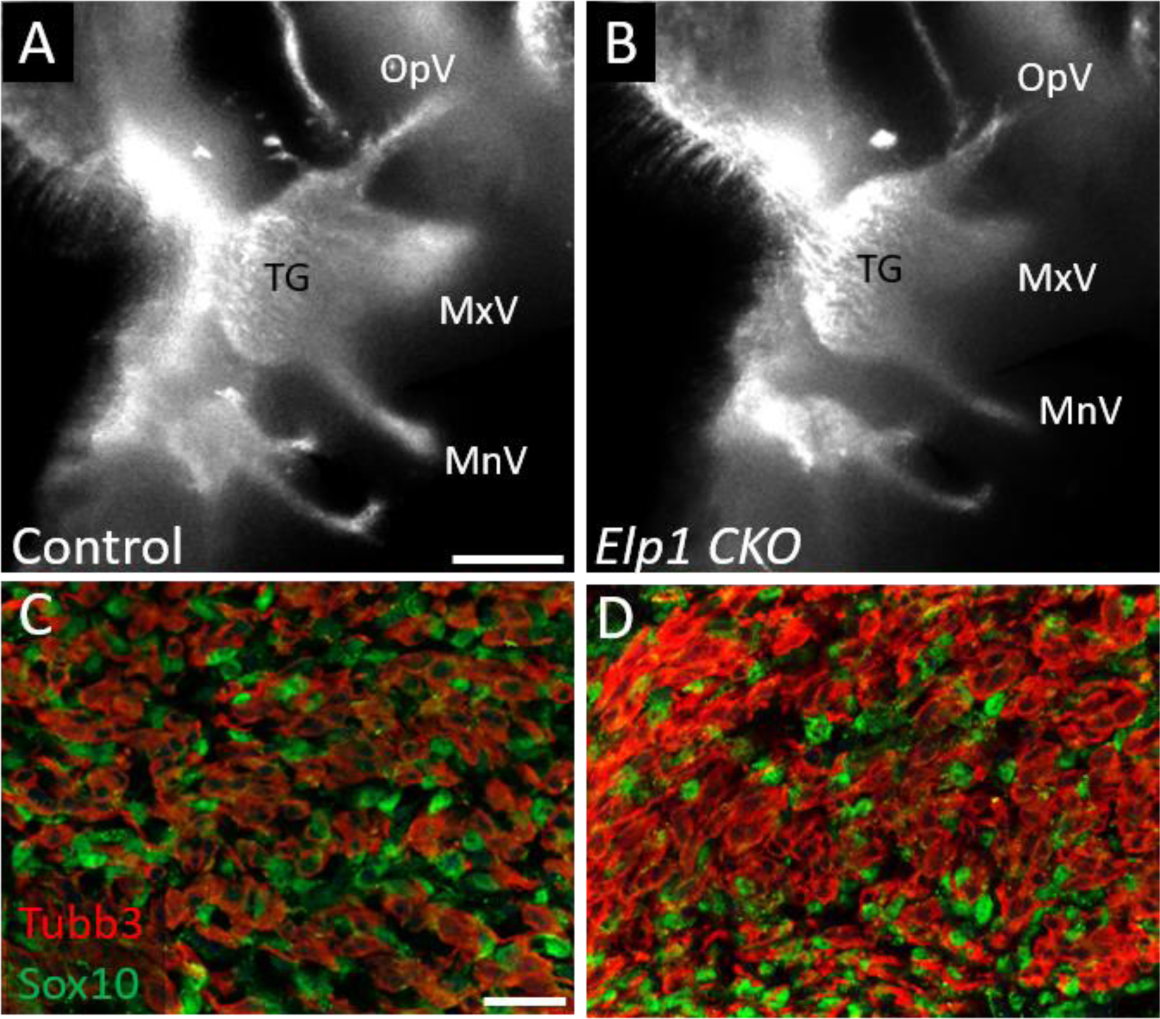
Neural crest cell migration and initial trigeminal ganglion formation appear normal in *Elp1 CKO* at E10.5. (**A-B**) Lateral view of the trigeminal ganglion of Control (**A**, N = 3) and *Elp1 CKO* (**B**, N = 3) after Tubb3 whole-mount immunohistochemistry (white). (**C-D**) Fluorescent immunohistochemistry on representative sections within the trigeminal ganglion from Control (**C**, N = or *Elp1 CKO* (**D**, N = 2) shows placodal neurons (Tubb3, red) and neural crest cells (Sox10, green). Scale bars: 400µm (**A**), also applies to **B**; 20µm (**C**), applies to **D**. Abbreviations: MnV – mandibular nerve, MxV – maxillary nerve, OpV – ophthalmic nerve, TG – trigeminal ganglion.

To capture the neurodevelopmental dynamics occurring in the trigeminal ganglion in control versus *Elp1 CKO*, we visualized intact trigeminal ganglia and associated nerves at intervals representing early (E11.5), middle (E12.5), and late (E13) stages of neural crest neurogenesis, using whole-mount Tubb3 immunohistochemistry. By E11.5, control and *Elp1 CKO* displayed robust outgrowth of axons from the trigeminal ganglion, which contains both neural crest- and placode-derived neurons at this stage (Fig. 4A, D). Although the ophthalmic, maxillary, and mandibular nerves were distinct in control and *Elp1 CKO*, axons in *Elp1 CKO* nerves appeared less organized. In the maxillary nerve, *Elp1 CKO* axons were observed straying from established axon bundles into the surrounding mesenchyme or crossing to adjacent fascicles (Fig. 4B, E). Axons of the mandibular nerve in *Elp1 CKO* also traveled away from the established nerve without direction, compared to the compact mandibular nerve observed in controls (Fig. 4C, F). By E12.5, all subdivisions of the ophthalmic nerve were severely deformed in *Elp1 CKO*, yet there were nuanced changes depending on the specific nerve (Fig. 5A, D). For example, the developing frontal nerve was present in both control and *Elp1 CKO*; however, the branching complexity of the nerve was decreased in *Elp1 CKO* (Fig. 5B, E), as quantified by a modified Sholl analysis (Fig. 5G, H, Sholl 1953). Proximal to the ganglion, stray axons were also observed growing away from established ophthalmic branches (Fig. 5E). While the rostral extension of the ophthalmic nerve was present in both control and *Elp1 CKO*, the medial and lateral nasal nerves that extend from this branch were only established in control embryos at this stage (Fig. 5C, F). Interestingly, though there was no obvious difference in the size of the central nerve root that contains axons projecting to the brainstem at E11.5 (Fig. 4A, D), the nerve root was smaller in *Elp1 CKO* by E12.5, indicating the loss of some central axons during this developmental time frame (Fig. 5A, D).

**Figure 4.**
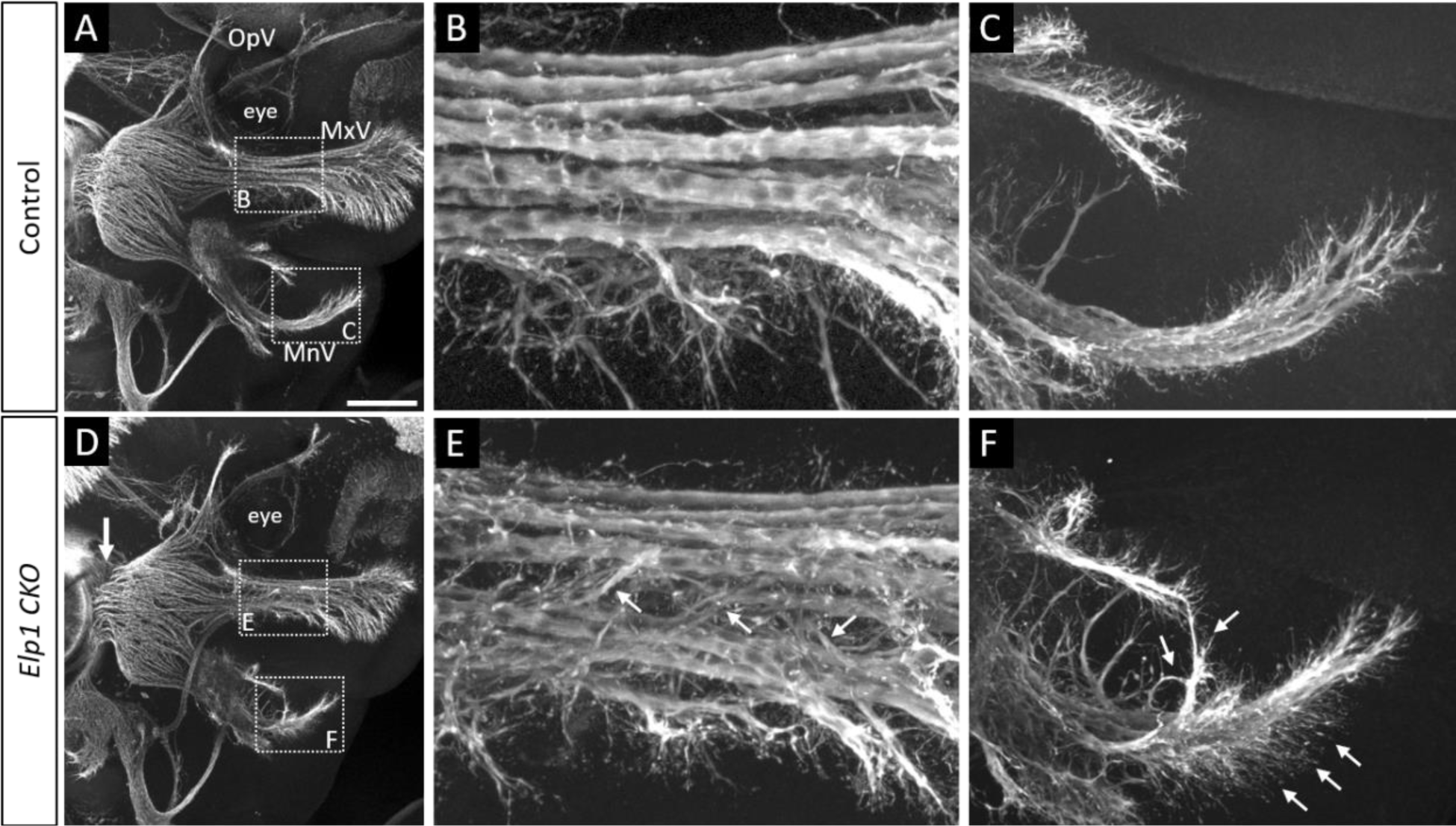
Progressive trigeminal nerve abnormalities arise in *Elp1 CKO* starting at E11.5. (**A-F**) Maximum intensity projections of confocal Z-stacks through Control (**A-C**, N = 3) or *Elp1 CKO* (**D-F**, N = 4), which were processed for whole-mount immunohistochemistry to detect Tubb3 (white), followed by tissue clearing. (**B**, **C**, **E**, **F**) Higher magnification images of boxed regions shown in **A** and **D**. Arrows indicate less compact central root (**D**) and disorganized axons (**E, F**) in *Elp1 CKO*. Scale bar: 200µm (**A**), applies to **B**; also applies to **B, C, E, F** as 50µm. Abbreviations: MxV – maxillary nerve, MnV – mandibular nerve, OpV – ophthalmic nerve.

**Figure 5.**
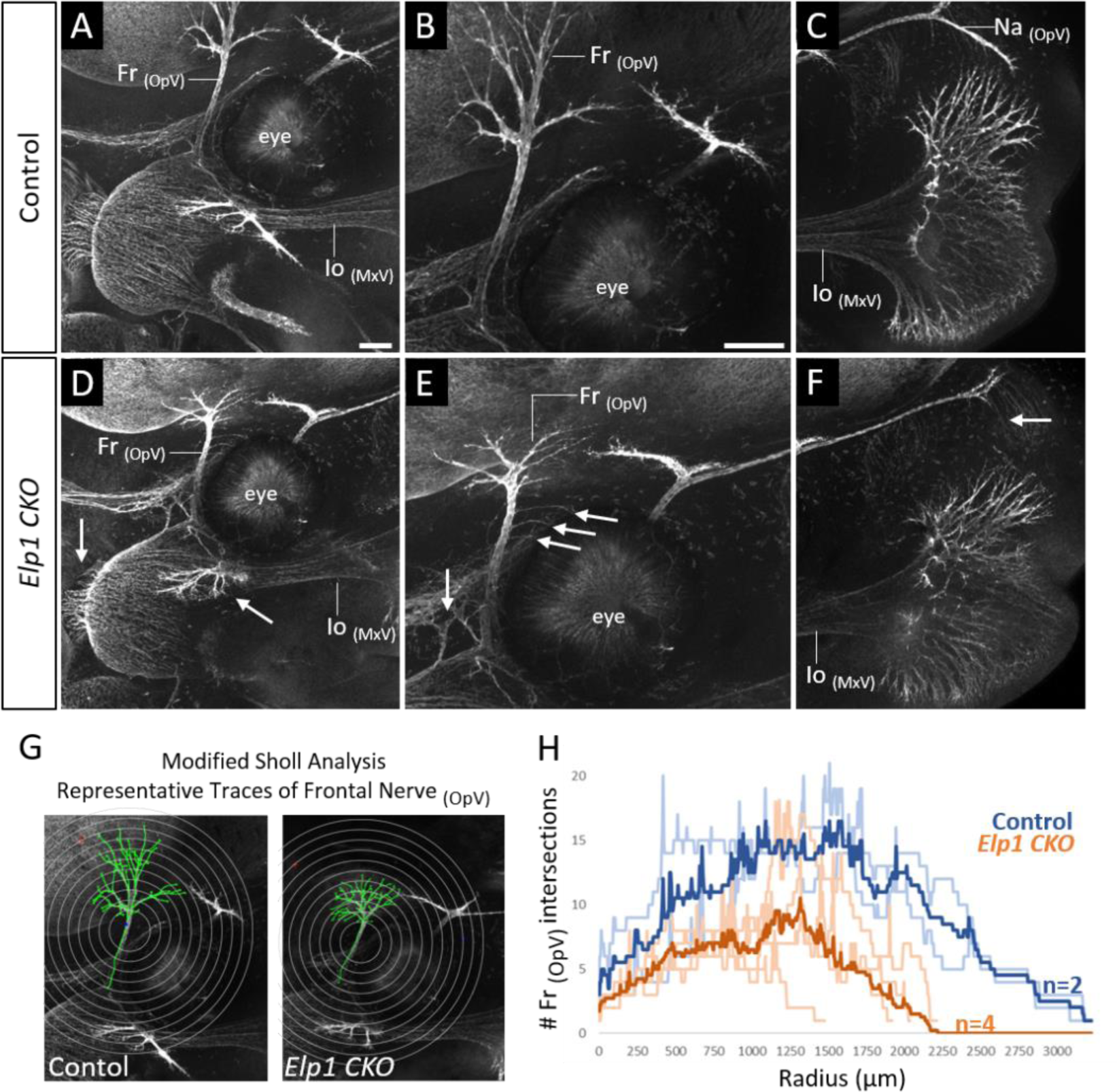
Trigeminal nerve branches are less complex or absent in *Elp1 CKO* at E12.5. (**A-F**) Maximum intensity projections of confocal Z-stacks through Control (**A-C**, N = 2) or *Elp1 CKO* (**D-F**, N = 4), which were processed for whole-mount immunohistochemistry to detect Tubb3 (white), followed by tissue clearing. (**B**, **C**, **E**, **F**) Higher magnification images of the frontal nerve division of the ophthalmic trigeminal branch around the eye (**B**, **E**) or the infraorbital nerve division of the maxillary branch at the developing whisker pad (**C**, **F**). Arrows indicate small central root and branch of ophthalmic nerve (**D**), disorganized axons (**E**), and the absence of the nasal nerve (ophthalmic division, **F**) in *Elp1 CKO*. (**G**) Diagram explaining Modified Sholl analysis, with concentric circles of increasing radii overlayed on representative traces (green) of the frontal nerve in Control (left) or *Elp1 CKO* (right). (**H**) Modified Sholl Analysis to quantify complexity of frontal nerve. Individual distributions are plotted in light blue (Control, N = 2) and light orange (*Elp1 CKO*, N = 4), while group averages are plotted in dark blue (Control) and dark orange (*Elp1 CKO*). Scale bar: 200µm (**A**), applies to **D**; 200µm (**B**), applies to **C, E, F**. Abbreviations: Fr - frontal nerve, Io – infraorbital nerve, MxV – maxillary nerve, OpV – ophthalmic nerve, TG – trigeminal ganglion.

Toward the end of the trigeminal ganglion neurogenic period, deficits in nerve growth and branching were even more pronounced in *Elp1 CKO*. The frontal nerve (ophthalmic) in *Elp1 CKO* was less complex at E13 than at E12.5, suggesting retraction and/or dying of some *Elp1 CKO* ophthalmic axons after failure to properly innervate the eye region (Fig. 6A, B). The lateral and medial nasal nerves (ophthalmic) were still absent in *Elp1 CKO* yet clearly established in controls (Fig. 6C, D). That the nasal nerves were not observed in *Elp1 CKOs* at any stage indicates some ophthalmic axons never reached their distal target site. Additionally, the infraorbital nerve (maxillary), which innervates the whisker pad, was less elaborate in *Elp1 CKO* compared to controls at E13 (Fig. 6C, D). Together, these results provide a dynamic summary of the requirements for Elp1 in axonal pathfinding and target innervation in trigeminal ganglion neurons. Importantly, our data suggest that nerve defects caused by loss of Elp1 can differ, even within the same ganglion, based on neuronal identity and/or specific target region.

**Figure 6.**
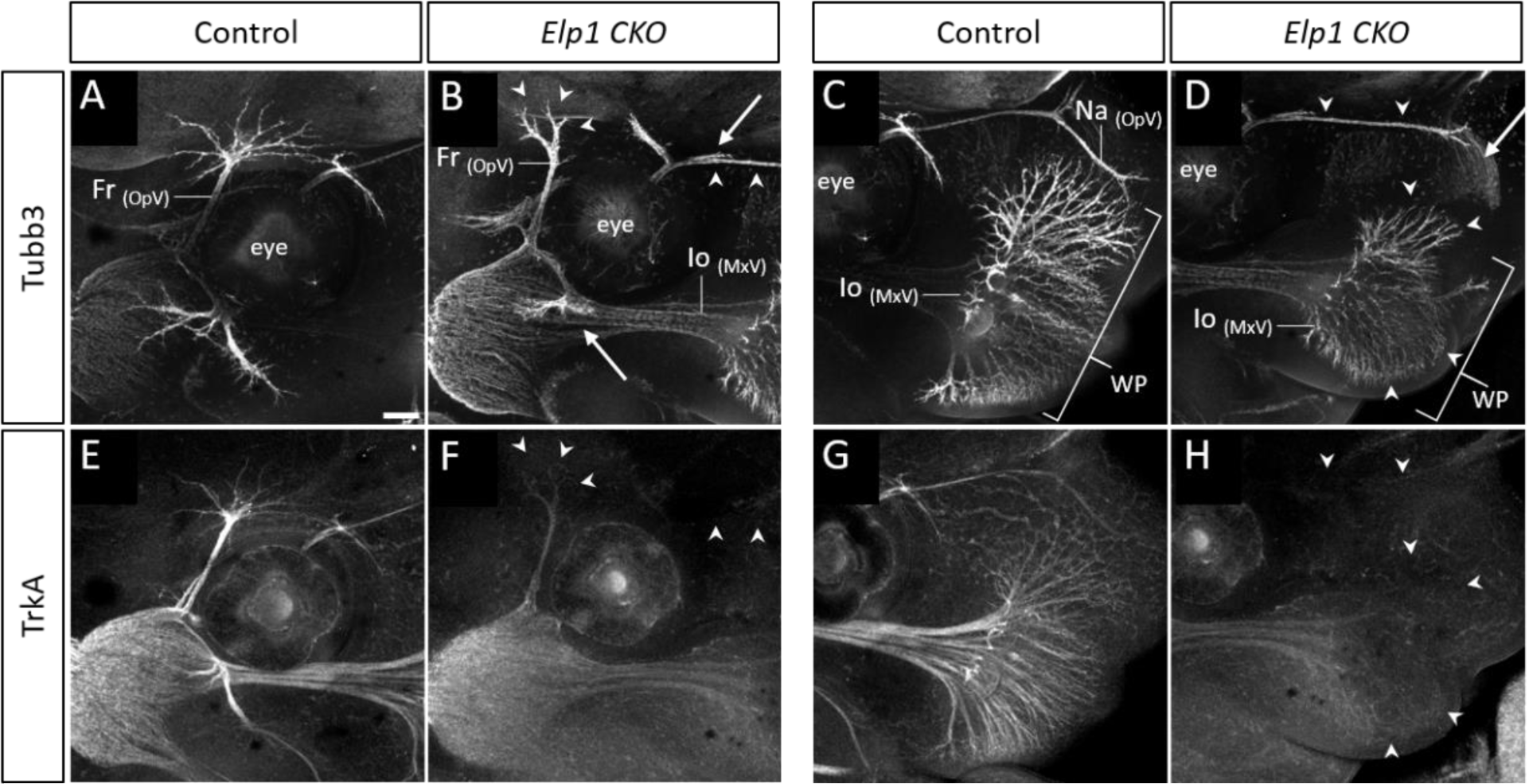
TrkA expression is decreased with persistent innervation defects in *Elp1 CKO* at E13. (**A-H**) Maximum intensity projections of confocal Z-stacks through Control (**A**, **C, E, G**, N = 2) or *Elp1 CKO* (**B**, **D**, **F, H**, N = 3), which were processed for whole-mount immunohistochemistry to detect Tubb3 (**A-D**) and TrkA (**E-H**), followed by tissue clearing. Arrows indicate regions where nerves are absent or severely diminished in *Elp1 CKO* (**B**, **D**), while arrowheads point to areas with Tubb3-positive nerves but undetectable TrkA expression (**B**, **D, F, H**). Scale bar: 200µm (**A**) and applies to all images. Abbreviations: Fr - frontal nerve, Io – infraorbital nerve, MxV – maxillary nerve, OpV – ophthalmic nerve, TG – trigeminal ganglion.

### Decreased TrkA correlates with increased cell death and loss of nociceptive neurons in *Elp1 CKO* trigeminal ganglia

Given the finding that trigeminal ganglion target innervation is disrupted in *Elp1 CKO*, combined with published studies from sympathetic neurons suggesting Elp1 is required for TrkA signaling (Li, Gruner, and Tourtellotte 2020; Jackson et al. 2014; Naftelberg et al. 2016), the distribution of TrkA expression was also examined via whole-mount immunohistochemistry at E13. In control embryos, TrkA signal was strong throughout the trigeminal ganglion and along all three major nerve divisions (Fig. 6E, G). In the frontal (ophthalmic) and infraorbital (maxillary) nerves, TrkA was expressed ubiquitously, including in the smallest, most distal branches (Fig. 6E, G). In *Elp1 CKO*, TrkA was detected, but weaker, in the trigeminal ganglion and in the core of the ophthalmic and maxillary branches (Fig. 6F). Strikingly, TrkA signal was drastically reduced in more distal nerve branches at E13, including the frontal nerve endings, the rostral ophthalmic projection, and the infraorbital axons of the whisker pad (Fig. 6F, H). Moreover, there was significant loss of TrkA in the central root and brainstem axon tracts, indicating effects on both central and peripheral projections at this stage (Figure 6-figure supplement 1). Therefore, target innervation defects in *Elp1 CKO* trigeminal ganglion neurons are associated with a loss of TrkA in the ganglion and nerves. These results are particularly intriguing given that TrkA neurons typically function in nociception, and FD patients experience a reduction or loss in the ability to sense facial pain and temperature.

**Figure 6–figure supplement 1.**
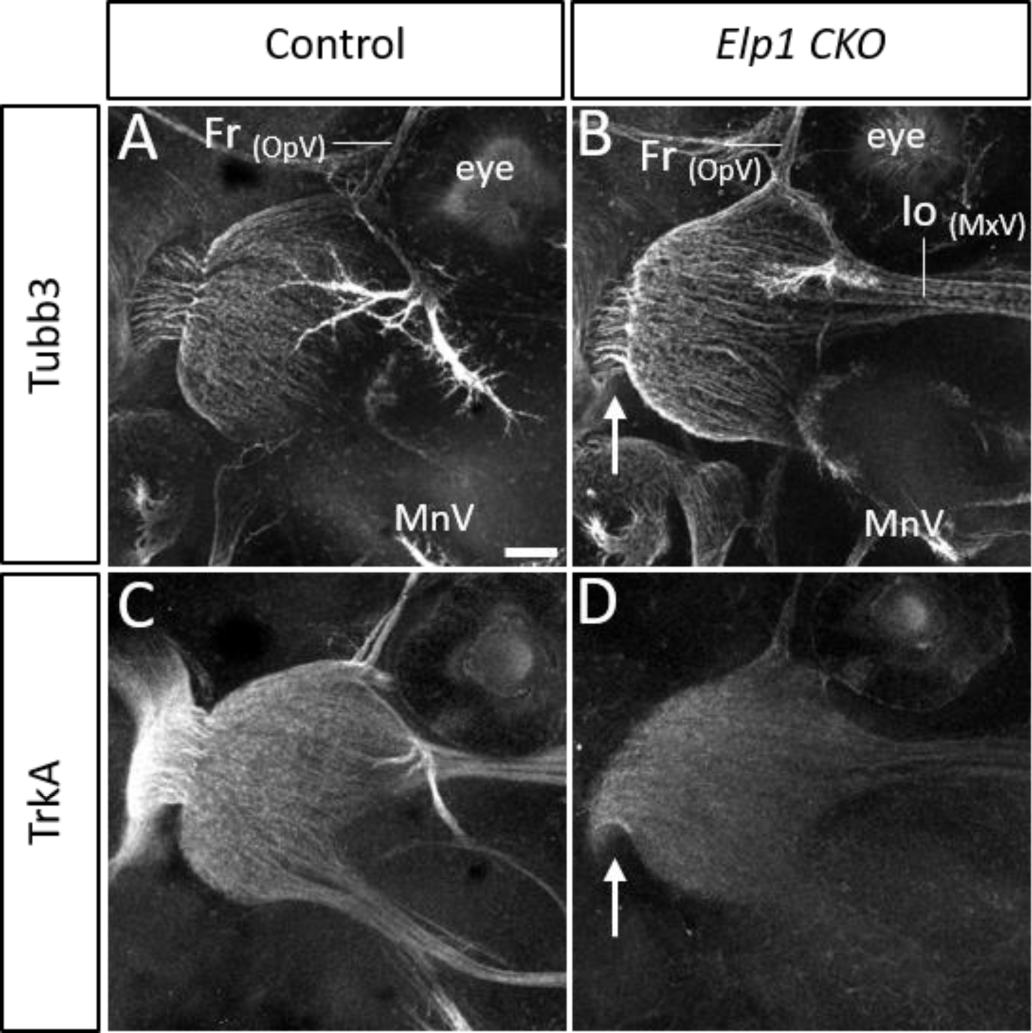
TrkA expression is decreased in the central nerve root of the trigeminal ganglion in *Elp1 CKO* at E13. (**A-D**) Maximum intensity projections of confocal Z-stacks through Control (**A**, **C,** N = 2) or *Elp1 CKO* (**B**, **D**, N = 3), which were processed for whole-mount immunohistochemistry to detect Tubb3 (**A-B**) and TrkA (**C-D**), followed by tissue clearing. Arrows point to central root in *Elp1 CKO* (**B**, **D**). Scale bar: 200µm (**A**) and applies to all images. Abbreviations: Fr - frontal nerve, Io – infraorbital nerve, MnV – mandibular nerve, MxV – maxillary nerve, OpV – ophthalmic nerve.

Next, we determined whether TrkA neurons were devoid of Elp1 protein in *Elp1 CKO* trigeminal ganglia. To this end, co-labeling for Elp1 and TrkA was performed via immunohistochemistry on control and *Elp1 CKO* tissue at E12.5. In control trigeminal ganglia, Elp1 was present in all neurons, and therefore co-labeled TrkA neurons, especially in axons (Fig. 7-figure supplement 1). In *Elp1 CKO* trigeminal ganglia, most TrkA neurons and axons did not express Elp1 (Fig. 7-figure supplement 1), while the remaining Elp1-positive neurons had large cell bodies characteristic of TrkB/C neurons (d’Amico-Martel and Noden 1980; D’Amico-Martel and Noden 1983; E. J. Huang, Zang, et al. 1999). To evaluate whether the loss of TrkA could be explained by neuronal cell death, TUNEL assays were also performed at E12.5, with co-labeling for TrkA. Compared to control embryos, which had occasional, scattered TUNEL-positive cells at this stage (Fig. 7A-C), *Elp1 CKO* had a dramatic increase in the number of TUNEL-positive cells distributed throughout the entire trigeminal ganglion, many of which also expressed TrkA (Fig. 7D-I). To determine whether this increased cell death resulted in the loss of specific neuronal subtypes, TrkA, TrkB, and TrkC neuron cell bodies were quantified in E13.5 trigeminal ganglia. While no differences were observed in the number of TrkB or TrkC neurons at this stage, TrkA neurons were reduced by 23 percent in *Elp1 CKO* trigeminal ganglia compared to controls (Fig. 7J, Control N=4; *Elp1 CKO* N=4; p = 0.016, unpaired t-test corrected for multiple comparisons with Holm-Sidak method). Collectively, these findings suggest that deletion of Elp1 from the neural crest lineage leads to detrimental innervation defects in the head, and the selective loss of TrkA neurons, as a result of widespread cell death within the trigeminal ganglion.

**Figure 7.**
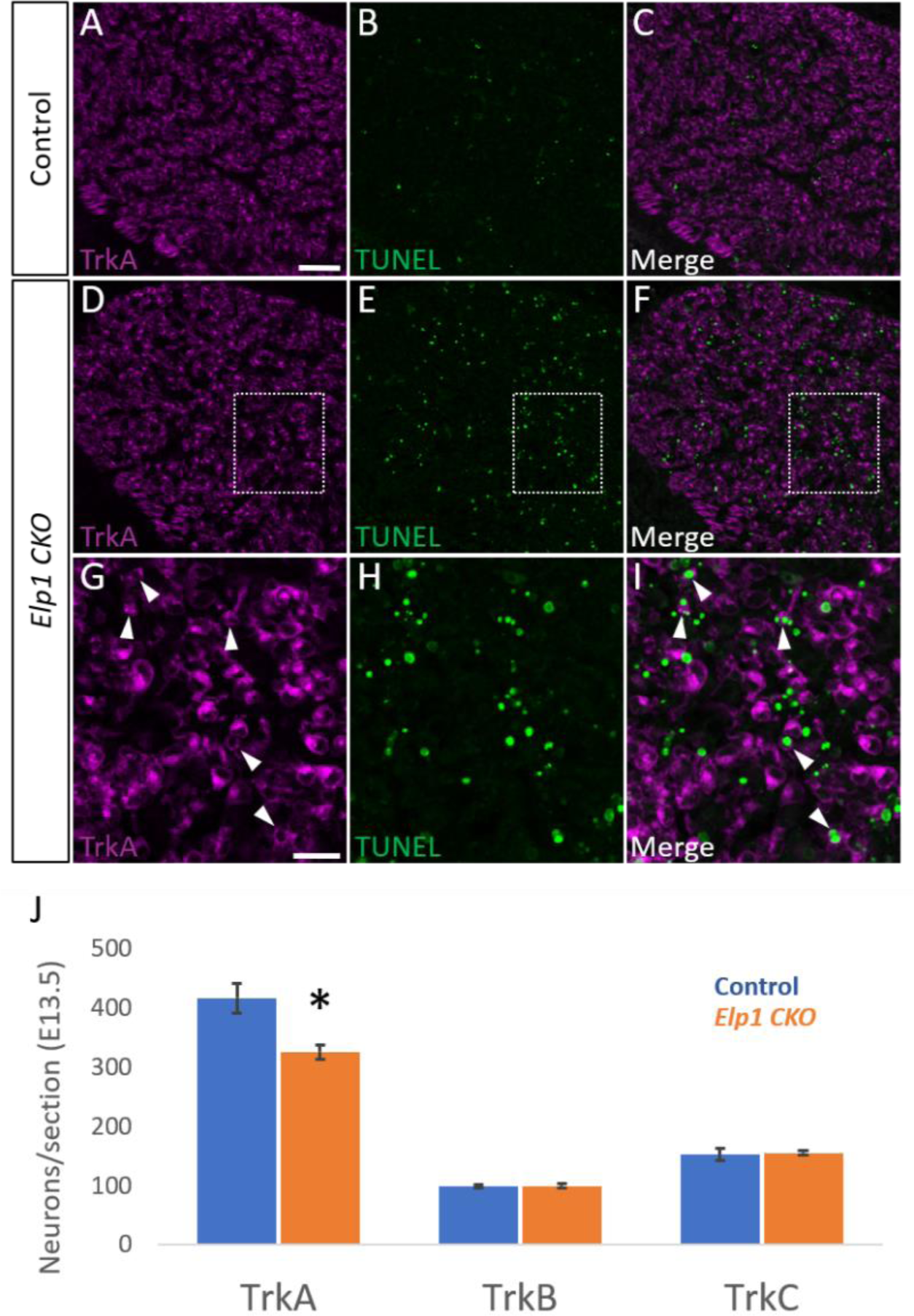
Loss of Elp1 from neural crest derivatives results in increased cell death and loss of TrkA neurons in the trigeminal ganglia. (**A-I**) Fluorescent immunohistochemistry on representative horizontal sections from E12.5 Control (**A-C**, N = 3) or *Elp1 CKO* (**D-I**, N = 3) revealing expression of TrkA (purple) with TUNEL staining (green). (**G-I**) Higher magnification of boxed regions in **D-F**. Arrowheads point to TrkA neurons that are TUNEL- positive (**G-I**). (**J**) Quantification of TrkA-, TrkB-, and TrkC-expressing neurons at E13.5 in Control (blue, N = 3) and *Elp1 CKO* (orange, N = 3) trigeminal ganglia. Values are mean ± SEM. *p = 0.016, unpaired t-test adjusted for multiple comparisons using Holm-Sidak method. Scale bars: 50µm (**A**), applies to **B-F**; 20µm (**G**), applies to **H-I**.

**Figure 7–figure supplement.**
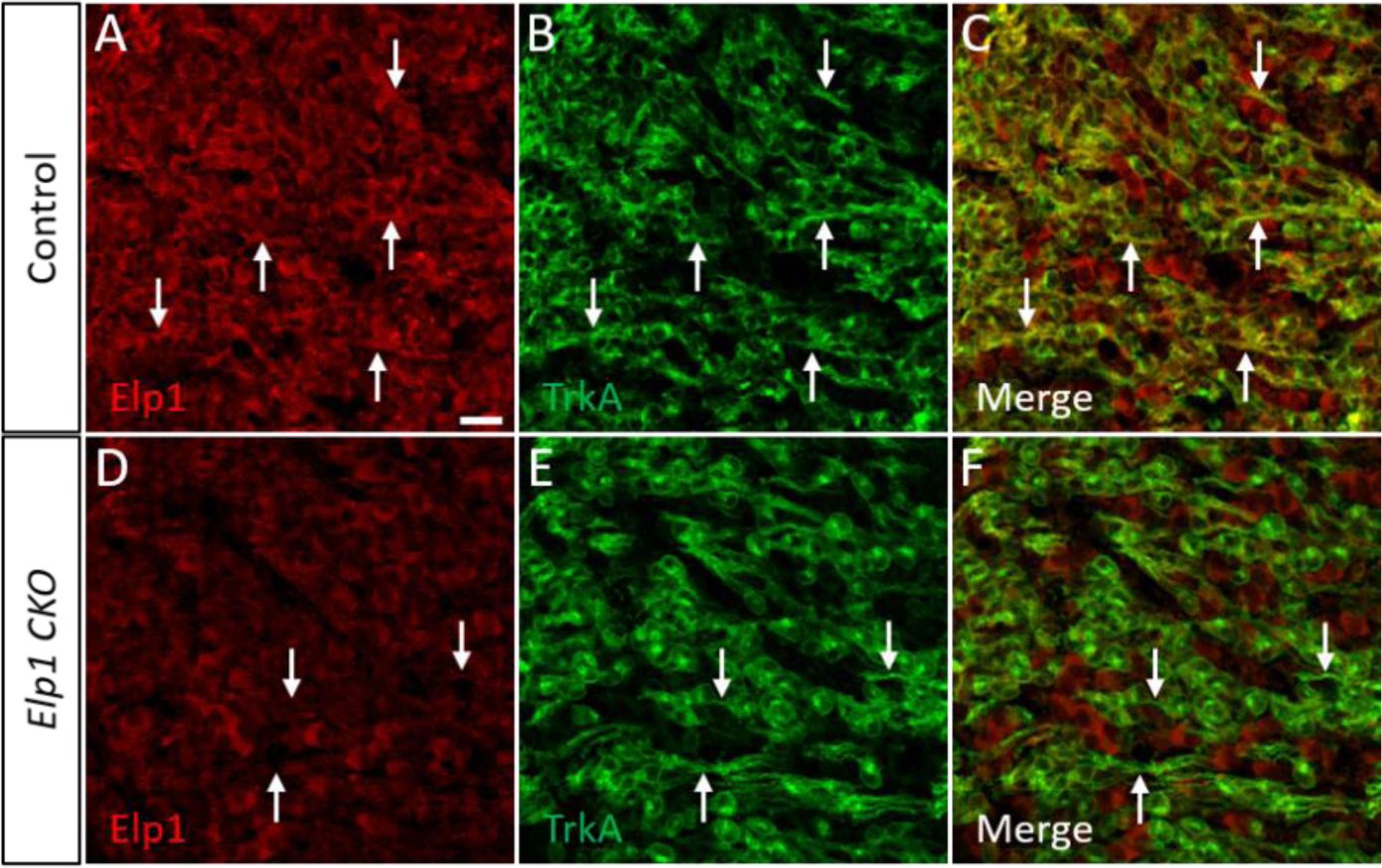
Loss of Elp1 from TrkA neurons in *Elp1 CKO* trigeminal ganglia. (**A-F**) Fluorescent immunohistochemistry on representative horizontal sections from E12.5 Control (**A-C**, N = 4) or *Elp1 CKO* (**D-F**, N = 4) mouse embryos demonstrating expression of Elp1 (red) and TrkA (green). Arrows point to TrkA-positive axons. Scale bar: 20µm (**A**) and applies to all images.

### *Elp1 CKO* provides evidence for the developmental origins of Trk-expressing neuronal subtypes in the trigeminal ganglion

Given the dual embryonic origin of the trigeminal ganglion, and our conditional knockout of *Elp1* from neural crest but not placode-derived tissues, we inquired about the ontogeny of Trk-expressing subpopulations to ask whether our findings could be due to interactions between neural crest and placode cells during early trigeminal ganglion development. First, we first explored the dynamics of normal TrkA neurogenesis in the trigeminal ganglion. Consistent with reports that TrkB and TrkC neurons are born first in the trigeminal ganglion (E9.5-11) followed by TrkA neurons (E11.5-13.5) (E. J. Huang, Wilkinson, et al. 1999; E. J. Huang, Zang, et al. 1999), immunohistochemistry on trigeminal ganglion sections revealed that all three Trk subtypes are present at E11.5 (Fig. 8A-F). At this stage, TrkC neurons comprise the largest proportion of neurons, followed closely by TrkA, with TrkB neurons representing the smallest proportion of trigeminal ganglion neurons (Fig. 8A-C). In accordance with previous findings (E. J. Huang, Zang, et al. 1999), the number of TrkB and TrkC neurons did not appear to increase between E11.5 and E12.5, but there was a robust increase in the number of TrkA neurons, such that TrkA neurons became the vast majority of trigeminal ganglion neurons by E12.5 (Fig. 8G-I). Together with our findings that Elp1 deletion from neural crest specifically targets TrkA neurons, and the fact that placode-derived neurons differentiate earlier than neural crest-derived neurons, we deduced that TrkA neurons in the trigeminal ganglion must be primarily neural crest-derived, while TrkB and TrkC neurons are placode-derived – a concept that has been alluded to in the literature, but not explicitly demonstrated.

**Figure 8.**
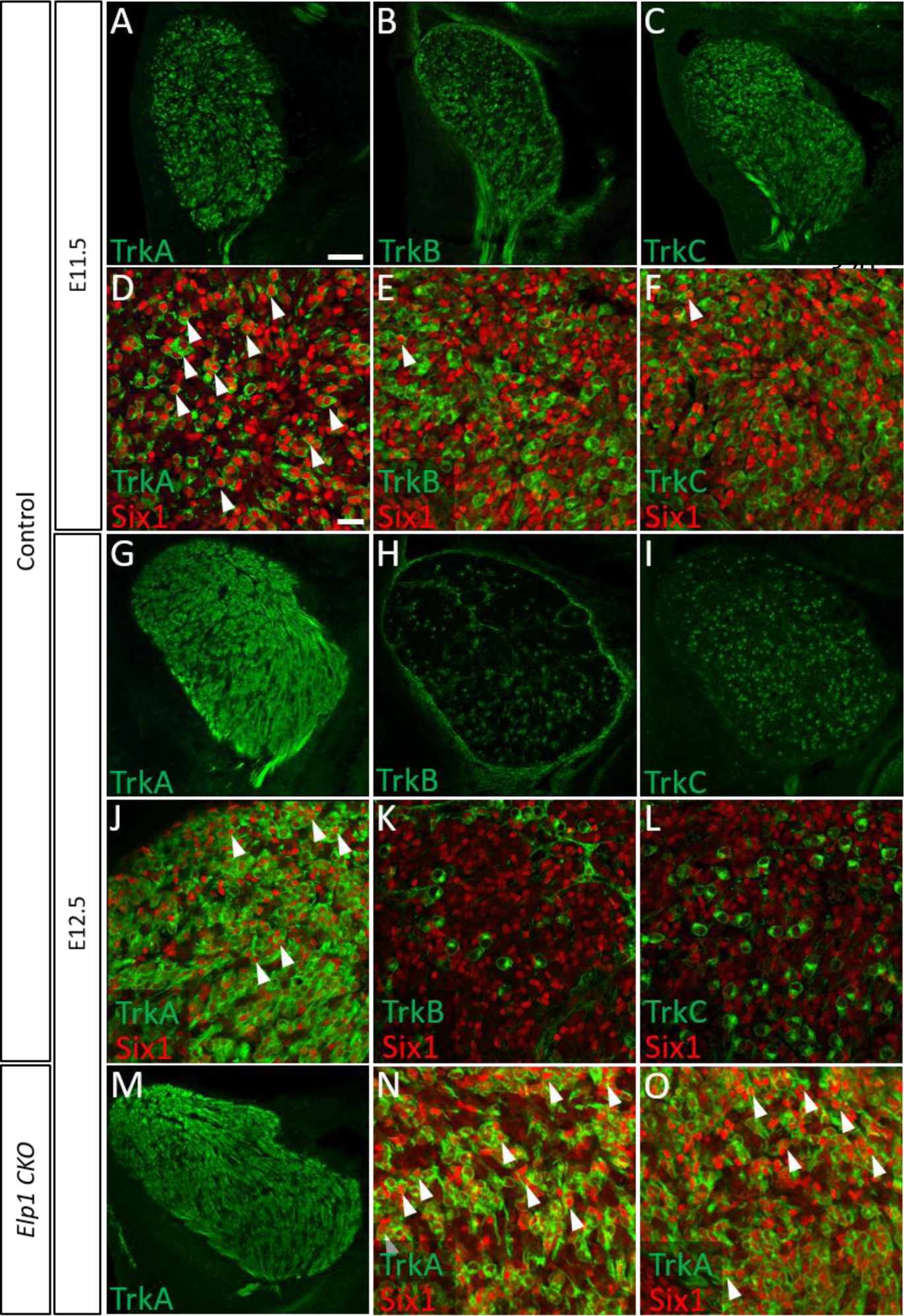
Dynamic expression of Six1 and Trk receptors occurs during trigeminal ganglion neurogenesis. (**A-O**) Fluorescent immunohistochemistry on representative horizontal sections from E11.5 (**A-F**) and E12.5 (**G-O**) Control (**A-L**, N=2 each) or *Elp1 CKO* (**M-O**, N=2) mouse embryos demonstrating expression of TrkA (**A, D, G, J, M-O**, green), TrkB (**B, E, H, K**, green), TrkC (**C, F, I, L**, green), and Six1 (**D-F, J-L**, **N-O**, red). Arrowheads point to neurons that co-express Six1 with TrkA (**D, J, N, O**), TrkB (**E**), or TrkC (**F**). Scale bars: 100µm (**A**), applies to **B-C, G-I, M**; 20µm (**D**), applies to **E-F, J-L, N-O**.

We aimed to validate these assumptions by co-labeling for the transcription factor Six1, which has historically been used as a marker of placode-derived neurons in the trigeminal ganglion (Karpinski et al. 2016; Moody and Lamantia 2015). Unexpectedly, we observed a large population of Six1-positive cells at E11.5, but these cells rarely expressed TrkB or TrkC. Instead, most of the Six1-positive cells at E11.5 expressed TrkA (presumptively neural crest-derived) or were devoid of any Trk receptor (Fig. 8D-F). At E12.5, the few neurons that strongly expressed Six1 also expressed TrkA, but never TrkB or TrkC (Fig. 8J-L). Given the timing of TrkA (E11.5-13.5) versus TrkB/TrkC (E9-11) neurogenesis, these data suggest Six1 is a marker of recently differentiated neurons in the trigeminal ganglion, rather than an exclusive marker of the placodal lineage. This claim is consistent with reports of Six1 expression in placode-derived neurons at E10.5 (Karpinski et al. 2016), and with our findings of Six1 expression in neural crest-derived TrkA neurons at E11.5-12.5. This conclusion is further supported by evidence that Six1 is transiently expressed in all newly differentiated, neural crest-derived neurons of the dorsal root ganglia (Yajima et al. 2014). To our knowledge, Six1 expression had not been examined in the trigeminal ganglion past E10.5, when the vast majority of neurons are still only placode-derived, thus explaining the gap in knowledge. With this information in hand, TrkA and Six1 expression was also examined in *Elp1 CKO* trigeminal ganglia. Six1- and TrkA-double positive neurons were still abundant in *Elp1 CKO* trigeminal ganglia at E12.5 (Fig. 8M-O). Altogether, these results suggest that neural crest-specific deletion of Elp1 leads to selective effects on neural crest-derived TrkA nociceptive neurons during the examined stages of trigeminal ganglion development, and that these effects largely occur after neuronal differentiation, ultimately resulting in decreased innervation and survival of trigeminal nociceptors.

## Discussion

Since the discovery that mutations in *ELP1* cause FD, several roles for Elp1 in neuronal development, function, and degeneration have been revealed (Slaugenhaupt et al. 2001; Anderson et al. 2001; Dietrich and Dragatsis 2016; Lefcort et al. 2017). To date, studies have emphasized defects in dorsal root ganglion and sympathetic neurons, which relay somatosensory and viscerosensory information from the trunk and limbs to the central nervous system (George et al. 2013; Jackson et al. 2014; Hunnicutt et al. 2012; Abashidze et al. 2014; Goffena et al. 2018; Li, Gruner, and Tourtellotte 2020; Morini et al. 2016). However, many questions remain regarding the cellular and molecular mechanisms underlying FD phenotypes, particularly since Elp1 has been proposed to mediate several cytoplasmic (e.g., protein trafficking, α-tubulin acetylation, stress signaling, exocytosis, and tRNA modification and translation of codon-biased transcripts) and nuclear (i.e., transcription) functions (Dalwadi and Yip 2018; Dietrich and Dragatsis 2016; Lefcort et al. 2017). Furthermore, while Elp1 knockdown in zebrafish leads to some defects in the formation of the trunk neural crest-derived enteric ganglia (Cheng et al. 2015), there is a dearth of information about Elp1 in the cranial ganglia, even when clinical deficits observed in FD clearly implicate trigeminal sensory dysfunction (Mendoza-Santiesteban et al. 2017; Barlow 2009; Geltzer et al. 1964; Gutiérrez, Norcliffe-Kaufmann, and Kaufmann 2015; Palma et al. 2018; Won et al. 2019). Perhaps this is due to the complexity of cranial ganglion development; dorsal root and sympathetic ganglia neurons are strictly neural crest-derived, whereas cranial ganglia contain neurons derived from both neural crest cells and ectodermal placodes. The trigeminal ganglion is especially unique, with intermingled placode- and neural crest-derived neurons (Blentic et al. 2011; D’amico-Martel 1982; D’Amico-Martel and Noden 1983; Hamburger 1961; Saint-Jeannet and Moody 2014; Steventon, Mayor, and Streit 2014). In this study, we provide the first in-depth analyses of the morphological and cellular consequences of *Elp1* deletion from the neural crest lineage during trigeminal ganglion development. Moreover, our data reveal insights on the contributions of neural crest versus placode cells to specific neuron populations and nerve branches emanating from the trigeminal ganglion.

### Elp1 is enriched in trigeminal neurons during the course of neurogenesis and early innervation

We first examined Elp1 expression in the developing trigeminal ganglion using an *Elp1:LacZ* reporter mouse, in addition to immunohistochemistry on wildtype tissue (Fig. 1 and Fig. 1–figure supplement 1). While we observed diffuse β-galactosidase staining throughout the neural tube, the trigeminal ganglia, and some cranial mesenchyme cells (Fig. 1A-C), Elp1 protein expression was more discrete, with robust expression in the cytoplasm of differentiated neurons (Fig. 1D-P, Fig. 1–figure supplement 1). These results mirror previously reported patterns of Elp1 protein expression in differentiated trunk sensory and sympathetic neurons (Hunnicutt et al. 2012; Jackson et al. 2014; Abashidze et al. 2014). Notably, little to no Elp1 protein was detected in developing glial cells within the trigeminal ganglion or in glia that line the embryonic nerves at the stages examined (Fig. 1N-P, Fig. 1–figure supplement 1), which represent the period of normal trigeminal ganglion neurogenesis and early innervation (E. J. Huang, Zang, et al. 1999; A. Davies and Lumsden 1984; Wilkinson et al. 1996). While the anatomical distribution of Elp1 protein overlapped with β-galactosidase staining in *Elp1:LacZ* mice, the observation that Elp1 protein is more selectively expressed in neurons suggests that Elp1 expression may be controlled at the post-transcriptional or post-translational level, such that Elp1 protein is upregulated upon neuronal differentiation. Altogether, our findings of highly enriched Elp1 expression in axonal compartments indicate extranuclear functions for Elp1 in developing trigeminal ganglion neurons, in agreement with previous studies that localize Elp1 to axonal transport and synaptic vesicles in other neuronal systems (Abashidze et al. 2014; Naftelberg et al. 2016; Tourtellotte 2016; Li, Gruner, and Tourtellotte 2020).

### Elp1 is required for trigeminal ganglion nerve outgrowth and innervation of target tissues

Next, we examined trigeminal ganglion development and trigeminal nerve deficits associated with FD using an established mouse model in which *Elp1* is conditionally deleted from neural crest cells and their derivatives (George et al. 2013). We found that Elp1 is not necessary for the migration of cranial neural crest cells to the developing ganglion, as the trigeminal ganglia in *Elp1 CKO* at E10.5 possessed appropriately condensed placodal neurons and neural crest cells and the gross morphology of the trigeminal ganglion was similar to controls (Fig. 3). These findings complement previously published results showing Elp1 is not required for the migration of trunk neural crest cells to dorsal root or sympathetic ganglia (George et al., 2013; Jackson et al., 2014). Interestingly, knockdown of another Elongator complex member, Elp3, in *Xenopus* neural crest cells inhibits their migration, suggesting that these Elongator subunits may have distinct functions in the neural crest (Yang et al. 2016).

As development proceeds, however, progressive defects in trigeminal nerve outgrowth and innervation become apparent in *Elp1 CKO* mice. By collecting high-resolution images of the intact ganglion and nerves over several timepoints, we observed various axonal abnormalities that preceded this outcome. In the ophthalmic, maxillary, and mandibular divisions of the trigeminal nerve at E11.5, axons generally extended in the direction of their respective targets, but many deviated from established fascicles along the way, resulting in disorganized nerves with a “hairy” appearance and ectopic branches in some locations (Fig. 4). These data suggest Elp1 may be required for axonal fasciculation in embryonic nerves and/or that Elp1 may regulate axonal branching in some trigeminal sensory neurons. Neuronal Elp1 depletion has been reported to increase or decrease neurite branching in different contexts (Hunnicutt et al. 2012; Abashidze et al. 2014; Jackson et al. 2014; Ohlen et al. 2017). While these varying observations could be due to variations in developmental timing, they may also highlight genuine differences in Elp1 function, depending upon the identity and environment of individual neurons.

The notion of context-dependent functions for Elp1 is further supported by our observations of different neuronal subtype responses to the loss of Elp1 within the trigeminal ganglion. By E12.5, at least some axons overcame aforementioned pathfinding deficiencies to reach their targets (Fig. 5). In the ophthalmic division of *Elp1 CKO*, the frontal nerve initially extended branches around the eye (albeit less elaborately than in controls, Fig. 5B, E, H), but then retracted, resulting in an even less complex pattern of innervation by E13 (Fig. 6A, B, E, F). Therefore, at least some frontal nerve axons in *Elp1 CKO* are able to reach their target site but are not maintained. These results suggest that Elp1 has a role in trigeminal sensory axon maintenance and neuronal survival as seen in other neuronal populations (Jackson et al. 2014; Li, Gruner, and Tourtellotte 2020; Naftelberg et al. 2016; Abashidze et al. 2014). In contrast to the initial invasion and subsequent retraction observed in frontal nerve axons, the medial and lateral nasal nerves (also ophthalmic division) never formed in the *Elp1 CKO* (Fig. 5C, F; Fig. 6C, D, G, H), indicating nasal nerve axons failed to navigate to their target destinations. Perhaps in these neurons, Elp1 is required for axons to respond to long-distance guidance cues. Further studies into the unique molecular, morphological, and functional identities of different trigeminal ganglion subpopulations are needed to understand the requirement for Elp1 in these different trigeminal neuronal subtypes.

### Elp1 is required for survival of TrkA neurons

Our data demonstrate that Elp1 is required for axon maintenance in some trigeminal ganglion neurons, indicating a potential role in neurotrophic signaling. Therefore, we interrogated the expression of Trk receptors in *Elp1 CKO* and control embryos. We found that loss of Elp1 from neural crest cells impacts TrkA neurons in the trigeminal ganglion, which generally develop into small diameter nociceptive neurons that relay pain and temperature information (Mu et al. 1993; Genç, Ulupinar, and Erzurumlu 2005; Reichardt 2006). Our results complement previous studies that demonstrate Elp1 is required for the survival of TrkA neurons in sympathetic and dorsal root ganglia (George et al. 2013; Jackson et al. 2014; Naftelberg et al. 2016; Li, Gruner, and Tourtellotte 2020; Ohlen et al. 2017). We observed a striking decrease in TrkA expression in the distal branches of the ophthalmic, maxillary, and mandibular nerves, as well as decreased TrkA signal in the central nerve root and the ganglion, itself (Fig. 6, Fig. 6–figure supplement 1). This could be explained by changes in the neurogenesis of TrkA-positive neurons, reduction in TrkA expression or trafficking, retraction of TrkA-positive axons, and/or death of TrkA neurons. While reduced TrkA numbers in dorsal root ganglia have been partially attributed to early cell cycle exit and cell death of Pax3-positive TrkA progenitors due to reduced DNA repair (George et al. 2013, Goffena et al., 2018), we did not observe Elp1 expression in Pax3-positive cells in the trigeminal ganglion (Fig. 1–figure supplement 1), which have been demonstrated as glial, not neuronal, progenitors (Baker, Stark, and Bronner-Fraser 2002). Moreover, Sox10-positive glia and glial precursors were present and normally distributed in *Elp1 CKO*, indicating, qualitatively, the absence of any initial effects on these cell populations (Fig. 3C-D).

Furthermore, we observe the initial presence of newly differentiated (Six1- positive), TrkA-positive, Elp1-negative neurons in *Elp1 CKO* trigeminal ganglia (Fig. 8M- O and Fig. 7–figure supplement 1). Therefore, our findings are most consistent with the idea that loss of Elp1 has major effects after neuronal differentiation; however, our analyses do not definitively rule out changes in neurogenesis in *Elp1 CKO*. In *Elp1 CKO*, we also observe that the majority of distal Tubb3-positive trigeminal nerve branches lack TrkA, whereas TrkA is robustly expressed into the smallest visible branches of the ophthalmic, maxillary, and mandibular nerves in controls at E13 (Fig. 6). Given the observation of widespread cell death in *Elp1 CKO* trigeminal ganglia at E12.5 (Fig. 7A-I) and the specific loss of TrkA neurons by E13.5 (Fig. 7J), it is unlikely that the remaining nerve endings in *Elp1 CKO* at E13 are from TrkA neurons. Collectively, our findings suggest Elp1 is required for trigeminal ganglion axons to grow, properly invade target tissues, and receive neurotrophic feedback that would maintain projections and prevent apoptosis, which are consistent with reports that severe innervation defects and subsequent apoptosis account for the majority of post-mitotic TrkA neuron loss in the trigeminal ganglia (Jackson et al. 2014; Abashidze et al. 2014; Li, Gruner, and Tourtellotte 2020).

In sympathetic neurons, Elp1 is reported to modulate TrkA/NGF retrograde signaling by regulating the phosphorylation of TrkA receptors in signaling endosomes (Li, Gruner, and Tourtellotte 2020). Our findings neither confirm nor negate the possibility of a similar mechanism for Elp1 in some trigeminal ganglion neurons. For axons that successfully navigate to their intended targets, deficient TrkA retrograde signaling could result in dying axons and subsequent cell death. Indeed, the retraction of axons that we note in the frontal nerve could indicate an inability to receive and/or appropriately respond to target-derived neurotrophic support. The observation that TrkA expression is severely diminished in distal axons (Fig. 6) could potentially contradict this mechanism or others that implicate deficiencies in TrkA retrograde signaling, since the phosphorylation state of TrkA receptors and/or retrograde transport speed is secondary to the requirement of the presence of adequate TrkA-positive receptors on distal axon terminals (Li, Gruner, and Tourtellotte 2020; Naftelberg et al. 2016). However, examination of earlier timepoints may reveal the initial presence of TrkA in Elp1-deficient nerve endings, followed by a decrease over time due to failed innervation and cell death. Detailed spatio-temporal studies are needed to gain a better understanding of the dynamics of TrkA expression and trafficking in trigeminal ganglion neurons and the role of Elp1 in these processes. Alternatively, Elp1 may be required for axonal outgrowth through other mechanisms, including anterograde trafficking of critical receptors for growth or guidance factors and/or regulation of local protein synthesis, all of which are required within growing axons (Scott- Solomon and Kuruvilla 2018; Ascaño et al. 2009; Cioni et al. 2019; Korsak et al. 2016; Batista and Hengst 2016; Kang and Schuman 1996). It is important to remember that these functions of Elp1 could be direct or indirect via its essential role as part of the Elongator complex in modifying tRNAs (B. Huang, Johansson, and Byström 2005; B. Huang, Jian, and Byström 2008; Y.-T. Chen et al. 2009; C. Chen, Tuck, and Byström 2009; Karlsborn, Tükenmez, Mahmud, et al. 2014; Karlsborn, Tükenmez, Chen, et al. 2014; Goffena et al. 2018). Indeed, the dorsal root ganglia proteome in *Elp1 CKO* exhibits substantial changes (compared to control) across a wide range of cellular functions, including axon guidance and pathfinding (Goffena et al. 2018), but whether similar proteome alterations occur in the trigeminal ganglion remain to be explored. Collectively, these diverse functions for Elp1 could explain, in part, the varying outcomes we observe across the trigeminal nerves, given the complexities associated with different target tissues, long-range signals, and neurotrophic factors required by different trigeminal neuron subpopulations. Moreover, the dual origin of the trigeminal ganglion makes it difficult to know whether these phenotypes are due to intrinsic differences in neural crest- versus placode-derived neurons, which will be critical to parse out in future studies.

### Insights into neural crest versus placodal cell dynamics in the developing trigeminal ganglion

During the course of this study, we made several observations on the temporal sequence of events during neurogenesis and axon growth in the developing trigeminal ganglion. Although the ganglion contains a mixture of neural crest- and placode-derived neurons, which we demonstrate both express Elp1 (Fig. 1), only neural crest-derived cells were targeted for *Elp1* deletion in these experiments. Since the vast majority of trigeminal ganglion neurons are placode-derived at E10.5 (Karpinski et al. 2016), it was not surprising that Tubb3 whole-mount immunohistochemistry, and trigeminal ganglion development, were similar between *Elp1 CKOs* and controls at this stage (Fig. 3). Striking alterations in the trigeminal ganglion and nerves were only noted in *Elp1 CKO* embryos after E10.5, once neural crest cells have begun differentiating into neurons. Neurons within the trigeminal ganglion will eventually express a distinct Trk receptor, which is typically correlated with a specific sensory modality (Patapoutian and Reichardt 2001). In *Elp1 CKO* embryos, TrkA neurons are vulnerable to loss of Elp1, with no impact on TrkB and TrkC neurons at the examined developmental stages.

The sequence of neural crest- versus placode-derived neuron differentiation may also clarify the developmental origin of Trk receptor-expressing neuronal subpopulations in the trigeminal ganglion. Previous studies demonstrated that, in the mouse trigeminal ganglion, TrkB and TrkC neurons are born before E11, whereas TrkA neurons are born on/after E11 (E. J. Huang, Wilkinson, et al. 1999; E. J. Huang, Zang, et al. 1999). As this timing coincides with placode-derived neurogenesis (starts at E9) preceding neural crest-derived neurogenesis (starts after E10.5, Karpinski et al. 2016), we propose that TrkB and TrkC neurons are likely placode-derived while TrkA neurons are primarily neural crest-derived. This notion is supported by the selective loss of TrkA neurons in neural crest targeted-*Elp1 CKO* trigeminal ganglia, the absence of initial effects on TrkB and TrkC neurons (Fig. 7J), and our findings that Six1, a marker of recently differentiated sensory neurons, co-localizes to TrkA neurons, and not TrkB or TrkC neurons, at E11.5 and E12.5 (Fig. 8). While extensive studies have defined Trk receptor expression and the timing of neural crest- vs. placode-derived neuron differentiation in the mouse trigeminal ganglion (E. J. Huang, Wilkinson, et al. 1999; E. J. Huang, Zang, et al. 1999; Wilkinson et al. 1996; Karpinski et al. 2016), this *Elp1 CKO* mouse has serendipitously allowed us to deduce the cellular origin of Trk-expressing neuronal subtypes. Although no differences in the number of TrkB or TrkC neurons were observed in our study, it is possible that (presumptive placode-derived) TrkB or TrkC neurons will be indirectly affected later in development, in response to the neural crest-specific loss of Elp1, given the important reciprocal interactions between the two lineages (Shiau et al. 2008; Shiau and Bronner- Fraser 2009; Wu and Taneyhill 2019; Steventon, Mayor, and Streit 2014; Wu et al. 2014). It would be informative to selectively delete Elp1 from placode-derived neurons only, or in addition to neural crest derivatives, in order to better understand the effects of cell type- specific loss of Elp1 on trigeminal ganglion development. Unfortunately, there are no trigeminal placode-specific Cre drivers available at this time to answer those questions.

Remarkably, the pattern of nerve deficits in *Elp1 CKO* embryos provides indirect evidence that certain branches of trigeminal sensory nerves may be specifically neural crest- or placode-derived. For instance, innervation of the whisker pad is initiated by E11.5 in both controls and *Elp1 CKO* (Fig. 4A, D); however, the innervation field does not increase substantially in the *Elp1 CKO* (Fig. 5C, F; Fig. 6C, D), and surviving nerve terminals are TrkA-negative, whereas control embryos develop a more expansive, TrkA- abundant innervation pattern by E13.5 (Fig. 6). This suggests that placode-derived neurons may reach the whisker pad first, and act as scaffolds upon which later-differentiating TrkA axons can traverse. A similar phenomenon is observed in the frontal nerve of the ophthalmic division in *Elp1 CKO* embryos, where the main branch is established by E11.5 (Fig. 4) and maintained at E13, but further elaboration and innervation are not supported over developmental time (Fig. 5D, E, H; Fig. 6A, B). In contrast, some trigeminal nerve branches never form in the *Elp1 CKO*, including the medial and lateral nasal nerves of the ophthalmic division (Fig. 5F, Fig. 6D), begging the question of whether these nerves are exclusively comprised of axons from neural crest-derived neurons. More in-depth studies are required to understand the diversity of interactions between neural crest- versus placode-derived cells during trigeminal ganglion neurogenesis and innervation, and to understand the nuanced roles that Elp1 has in such a complex system.

FD is characterized by a wide range of sensory and autonomic phenotypes, including those that implicate the involvement of the cranial trigeminal ganglion and its associated nerves. Through studies in a conditional knockout mouse model that eliminates Elp1 from the neural crest lineage, we have uncovered distinct effects on the development of trigeminal sensory neurons, specifically those expressing TrkA that generally function in nociception. Our results introduce a mechanistic basis for the neuronal deficits that contribute to the loss of facial pain and temperature sensation experienced by FD patients.

## Materials & Methods

### Animal Husbandry

All animal care and use described herein was in accordance with federal and institutional guidelines and approved by Montana State University’s and University of Maryland’s IACUC, under protocols #2018-81 (MSU) and #R-MAR-20-15 (UMD). The generation of the *Elp1* conditional knockout mouse was described previously by Dr. Frances Lefcort (Montana State University), who generously provided the embryos for this work (George et al. 2013). All mice were created and maintained on a C57BL/6 background. *Elp1^fl/fl^* mice have *LoxP* sites flanking the coding region of the fourth exon of *Elp1* (previously *Ikbkap*). When the floxed region is excised via Cre recombinase, the resulting truncated *Elp1* transcript is eliminated from cells by nonsense-mediated decay. The role of Elp1 in Wnt1-expressing neural crest cells and derivatives was examined by crossing homozygous *Elp1^fl/fl^* mice with hemizygous *Wnt1-Cre^+/-^* mice (The Jackson Laboratory, stock no. 003829) to create Elp^fl/+^;Wnt1-Cre^+/-^ males, which were then crossed with Elp1^fl/fl^ females to generate *Elp1* conditional knockout mice (*Elp1^fl/fl^; Wnt1-Cre^+^*, abbreviated “*Elp1 CKO”* throughout). For all analyses, *Elp1 CKO* mice were compared to littermate controls (*Elp1^fl/+^; Wnt1-Cre^-^*) and at least 2 litters were examined per experiment. Genotyping was performed via PCR using the primer sequences listed below. For timed breeds, the day of the vaginal plug was considered embryonic day 0.5 (E0.5). Samples sizes were determined based on previously published studies using this mouse model or similar *Elp1* conditional knockouts mouse models (George et al. 2013; Jackson et al. 2014; Li, Gruner, and Tourtellotte 2020).

### Genotyping

Genomic DNA was extracted using the Extracta DNA Prep for PCR kit (Quantabio) according to manufacturer instructions. *Elp1* CKO and wildtype alleles were detected with the following primers: 5’-GCACCTTCACTCCTCAGCAT-3’ (forward) and 5’- AGTAGGGCCAGGAGAGAACC-3’ (reverse). The *Wnt1-Cre* allele was detected with the following primers: 5’-GCCAATCTATCTGTGACGGC-3’ (forward) and 5’- CCTCTATCGAACAAGCATGCG-3’ (reverse). PCR mixtures were prepared using DreamTaq Green PCR Master Mix (Thermo Fisher) according to the manufacturer’s protocol.

### Tissue Collection and Preparation

Timed pregnant females were euthanized via CO2 asphyxiation followed by cervical dislocation. Embryos were collected and placed in ice-cold 1X phosphate-buffered saline (1X PBS). A hindlimb bud was collected from each embryo for genotyping. Embryos were fixed via submersion and gentle shaking in 4% paraformaldehyde/1X PBS for between 20 minutes (E10.5) to 90 minutes (E13.5) at room temperature, then rinsed three times in 1X PBS for 20 minutes each. Fixed embryos were stored in 1X PBS with 0.02% sodium azide at 4°C until further analysis. For sectioning, embryos were rinsed twice with 1X PBS, then submerged in 15% sucrose (w/v) in 1X PBS at 4°C overnight, or until tissue sank, followed by submersion in 30% sucrose at 4°C until tissue sank. Embryos were first equilibrated in a 1:1 solution of 30% sucrose/1X PBS and Tissue-Tek Optimal Cutting Temperature compound (OCT, Fisher) for 2 hours at 4°C and then in 100% OCT at 4°C for 2 hours. Embryos were embedded in 100% OCT using liquid nitrogen vapor and stored at -80°C, followed by sectioning at 12µm on a cryostat (Leica) and collection of tissue sections on Superfrost Plus charged slides (VWR).

### Immunohistochemistry

#### Tissue Sections

A hydrophobic boundary was drawn around tissue sections using an ImmEdge Pen (Vector Labs). Tissue sections were rehydrated with 1X PBS for 5 minutes, then permeabilized with 1X PBS/0.5% Triton X-100 (Tx-100) for 5 minutes at room temperature. Tissue was blocked with 5% Bovine Serum Albumin (BSA, Fisher Scientific) (w/v) in PBS- Tx (1X PBS, 0.1% Tx-100) for approximately 1 hour at room temperature, then rinsed once in PBS-Tx. Primary antibodies were diluted in PBS-Tx plus 1% BSA and applied overnight at 4°C in a humidified chamber. Unbound primary antibodies were washed off with 4 PBS- Tx rinses for 5 minutes each at room temperature. Sections were then incubated with secondary antibodies, diluted in PBS-Tx plus 1% BSA, for 1 hour at room temperature in a humidified chamber. Sections were rinsed 3 times in PBS-Tx for 5 minutes each, followed by 2 rinses in 1X PBS for 5 minutes each, all at room temperature. Coverslips were mounted with DAPI Fluoromount-G Mounting Medium (Southern Biotech) and allowed to dry in the dark at room temperature overnight before imaging.

#### Whole-mount

Fixed embryos were rinsed twice with 1X PBS for 5 minutes per rinse, then dehydrated through a series of increasingly concentrated methanol (MeOH) washes (50:50, 80:20, 100:0 MeOH:PBS) for 30 minutes each at room temperature. Embryos were incubated in Dent’s Bleach (4:1:1 MeOH:DMSO:30% H2O2) for 6 hours at room temperature with gentle shaking, then rehydrated through a series of decreasingly concentrated methanol washes (50:50, 20:80, 0:100 MeOH:PBS) for 30 minutes each at room temperature. For blocking, embryos were incubated in antibody dilution solution (1X PBS, 0.1% Tx-100, 5% BSA) for at least 2 hours at room temperature. Next, embryos were incubated with fresh antibody dilution solution containing primary antibodies for 2 days (E10.5) up to 4 days (E13.5) at 4°C with gentle shaking. Embryos were washed 4 times for ∼1 hour each at room temperature with PBS-Tx, then incubated in fresh dilution solution with secondary antibodies for 1 day (E10.5) up to 3 days (E13.5) at 4°C with gentle shaking. Embryos were washed 3 times for ∼1 hour each at room temperature with PBS-Tx, followed by 2 washes with 1X PBS for 30 minutes each at room temperature. E10.5 embryos were imaged at this step, whereas older embryos were cleared before imaging, as described below.

### Antibodies

Primary antibodies used included the following: Elp1 (Sigma #SAB2701068, 1:500), β- tubulin III (Abcam #ab78078, 1:1000 for sections, 1:300 for whole-mount), Sox10 (R&D #AF2864, 1:200 or GeneTex #GTX128374, 1:500), TrkA (R&D #AF1056, 1:500 for sections, 1:200 for whole-mount), TrkB (R&D #AF1494, 1:300), TrkC (R&D #AF1404, 1:300), Six1 (Sigma #HPA001893, 1:500), Islet1 (DSHB #PCRP-ISL1-1A9, 1:500), Neuropilin2 (R&D cat. AF567, 1:500), and Pax3 (DSHB, “Pax3”, 1:200). All species/ isotype-specific, Alexafluor-conjugated secondary antibodies were purchased from Thermo Scientific and used at a dilution of 1:500 on sections, or 1:300 in whole-mount.

### FRUIT Clearing

After whole-mount immunohistochemistry, embryos were subjected to FRUIT clearing (Hou et al. 2015). Briefly, embryos were moved through a series of aqueous FRUIT buffers, containing 8M urea (Sigma), 0.5% (v/v) α-thioglycerol (TCI America), and increasing concentrations of fructose (Fisher). Embryos were incubated at room temperature with gentle rocking in 35% FRUIT for 6 hours, followed by 40% FRUIT overnight, 60% FRUIT for 8 hours, and 80% FRUIT overnight. Embryos were stored in 80% FRUIT at 4°C until imaging in this buffer.

### TUNEL Staining

TUNEL staining was performed on tissue sections after immunohistochemistry using the *In Situ* Cell Death Detection Kit, TMR Red (Roche) according to the manufacturer’s instructions. After washing off unbound secondary antibodies, slides were post-fixed with 4% PFA in 1X PBS for 5 minutes at room temperature, then washed twice with 1X PBS for 5 minutes each at room temperature. Sections were incubated with TUNEL reaction mixture for 60 minutes at 37°C in the dark, followed by 3 washes in 1X PBS for 5 minutes each at room temperature. Coverslips were mounted using DAPI Fluoromount-G Mounting Medium (Southern Biotech) and allowed to dry in the dark at room temperature overnight before imaging.

### Imaging

E10.5 embryos that underwent whole-mount β-galactosidase staining or fluorescent immunohistochemistry were imaged on a Zeiss SteREO Discovery V8 Pentafluor fluorescent microscope using AxioVision software (Zeiss). Embryos E11.5 and older that were processed for whole-mount immunohistochemistry were imaged in 80% FRUIT buffer on a Zeiss LSM 800 confocal microscope. Z-stacks were collected at 5µm intervals using 5X or 10X air objectives. Fluorescent immunohistochemistry on tissue sections was also visualized on the Zeiss confocal microscope using 10X and 20X air objectives, or the 63X oil objective. For all applications, laser power, gain, offset, and digital zoom were identical when imaging comparable regions of interest in Control versus *Elp1 CKO* embryos and the pinhole was set to 1 airy scan unit at all times. CZI files were processed in Zen software (Blue edition 2.0, Zeiss). For Z-stacks, CZI files were processed in ImageJ, where Maximum Intensity Projections were created using the Z-Project function in Hyperstack mode.

### Nerve Tracing & Sholl Analysis

Tracing of the frontal nerve was performed on TIFF images of maximum intensity Z-stack projections, described above, using the Simple Neurite Tracer plug-in in Image J (NIH). After tracing, the Sholl Analysis function within the Simple Neurite Tracer plug-in was used to quantify nerve branching complexity under the following settings: use standard axes, no normalization of intersections, and 10µm radius step size. The center point was set on the primary frontal nerve branch, just below the first branch point. Individual distributions and group average distributions were plotted together in Microsoft Excel (N = 2 Control, N = 4 *Elp1 CKO*, from two different litters).

### Quantification of Trk-expressing neurons

10X images were acquired that encompassed the entire trigeminal ganglion within each horizontal section, where tissue was co-labeled for Tubb3 and TrkA, TrkB, or TrkC. The boundary of the trigeminal ganglion was determined in each section by Tubb3 staining, which marked all neurons. Using the Cell Counter plug-in in ImageJ, each TrkA, B, or C neuron was counted within the pre-determined trigeminal ganglion boundary. Neurons were counted across 5 serial sections per animal, at the level of the maxillary nerve, and the average number of neurons per section was calculated and presented as mean ± standard error. The number of TrkA, B, or C neurons per section in *Elp1 CKO* was compared to Control using an unpaired t-test, adjusted for multiple comparisons using the Holm-Sidak method (Graphpad, N = 4 Control, N = 4 *Elp1 CKO* embryos, from two different litters).

## Acknowledgements

We thank Marta Chaverra for assistance with collecting embryos, Lynn George for guidance regarding genetics, Vickie Riojas for help with animal husbandry, and Jenn Lafrican for technical assistance.

